# MultiOmicsAgent: Guided extreme gradient-boosted decision trees-based approaches for biomarker-candidate discovery in multi-omics data

**DOI:** 10.1101/2024.07.24.604727

**Authors:** Jens Settelmeier, Sandra Goetze, Julia Boshart, Jianbo Fu, Sebastian N. Steiner, Martin Gesell, Peter J. Schüffler, Diyora Salimova, Patrick G. A. Pedrioli, Bernd Wollscheid

## Abstract

MultiOmicsAgent (MOAgent) is an innovative, Python based open-source tool for biomarker discovery, utilizing machine learning techniques specifically extreme gradient-boosted decision trees to process multi-omics data. With its cross-platform compatibility, user-oriented graphical interface and a well-documented API, MOAgent not only meets the needs of both coding professionals and those new to machine learning but also addresses common data analysis challenges like data incompleteness, class imbalances and data leakage between disjoint data splits. MOAgent’s guided data analysis strategy opens up data-driven insights from digitized clinical biospecimen cohorts and makes advanced data analysis accessible and reliable for a wide audience.

**Biographical Note:** Jens Settelmeier, Julia Boshart, Martin Gesell are Ph.D. candidates, Jianbo Fu, Sebastian N. Steiner are Post Doc candidates and Sandra Goetze, Patrick Pedrioli senior scientists at the Institute of Translational Medicine at Health Sciences and Technology department at ETH Zürich, Switzerland, within Professor Bernd Wollscheid’s research group who has been working in the fields of bioinformatics, clinical multi-omics with a focus on spatial cell surface proteomics.

Peter J. Schüffler is professor at the institute of Pathology at the TU Munich, Germany and has been working in the field of digital pathology and clinical multi-modal studies.

Diyora Salimova is junior professor at the department of Applied Mathematics at the Albert-Ludwigs-University of Freibug, Germany and has been working in the field of stochastic processes, approximation theory and machine learning related topics.

**Key Points:**

- MOAgent enables a guided biomarker-candidate discovery in multi-omics studies, providing a graphical interface and well-documented API.
- A user can run MOAgent on a personal computer without the requirement of coding a single line.
- MOAgent is a Python-based solution for biomarker-candidate discovery, using machine learning to analyze multi-omics data.
- MOAgent can address challenges like data incompleteness and class imbalances, ensuring reliable analysis.
- MOAgent makes advanced data analysis accessible, enhancing insights from clinical data.

## Introduction

The concept of a molecular digital twin, obtained through digitizing individual clinical biospecimens using multi-omics strategies, offers new opportunities to gain diagnostic insights into human health [1]. Next-generation sequencing-based transcriptomics, along with Mass Spectrometry (MS)-based proteomics and both MS and Nuclear Magnetic Resonance (NMR)-based metabolomics, are the de facto standard methodologies for the generation of quantitative data matrices representing the identity and abundances of thousands of transcripts, proteoforms, and metabolites across biological samples of interest [2–9]. Identifying the subset of molecular entities that strongly associate with biological classes of interest from these quantitative matrices is an essential first step in understanding the underlying class-specific molecular biology and defining putative biomarker signatures.

Over the past decade, the advancements in the development of computational tools for multi-omics analysis, particularly in enabling the detection of class-specific features, have been made by several researchers [10–15]. Despite these advances, multi-modal data analysis remains challenging. Analysts are required to efficiently manage incomplete data, complex combinations of multiple features, and false discovery rate (FDR) in the presence of limited replicates and large feature numbers. Indeed, (multi-)omic studies are often burdened by the curse of dimensionality and high rates of missing values [16]. This is often especially true in the context of analyzing data from clinical studies where sample cohorts of limited size and high heterogeneity can hinder the extraction of clinically relevant insights. Furthermore, interpreting the results from such analyses requires a strong understanding of the biological system under investigation. It is, therefore, desirable to make consistent multi-modal data analysis accessible to a wide scientific audience, notwithstanding coding or machine learning expertise. This can be achieved via user-friendly interfaces and protection from common pitfalls and challenges such as class imbalances, overfitting, generalization, and information leakage.

Here, we present MOAgent as a solution, an easy-to-use, open-source, cross-platform compatible, graphical user interface-driven application for selecting phenotypic features from quantitative (multi-)omics data using machine learning. MOAgent can directly handle molecular expression matrices - including proteomics, metabolomics, transcriptomics, as well as combinations thereof. The MOAgent-guided data analysis strategy is compatible with incomplete matrices and limited replicate studies.

## Materials and Methods

### Implementation and Features

The code for the MOAgent application comprises two Python packages that provide backend (https://github.com/Wollscheidlab/MOBiceps) and frontend (https://github.com/Wollscheidlab/MOAgent) functionalities. The former includes a number of functions centered around the selection and evaluation of phenotype-relevant features from quantitative omic-data and exposes a well-defined API for programmatic control by a coding-competent audience. The latter provides an intuitive GUI and streamlines access to these functions for users who might lack coding or machine learning expertise. In addition to GitHub and pip distributions, MOAgent is also available as a downloadable virtual machine.

The core functionality of MOAgent can be accessed via the “RFE++” section of the GUI (Figure F1). From this panel, input files and analysis parameters can be specified before starting the workflow. The newly implemented main algorithm used in the selection of phenotypic features is based on an improved version of the recursive feature elimination approach by Guyon et al. [17] we first implemented and used in [18,19] hereby referred to as RFE++. Specifically, we enhanced the compatibility for data with missing values (incompleteness), imbalanced class distributions, small datasets, and multi-omic studies.

**Figure F1:**
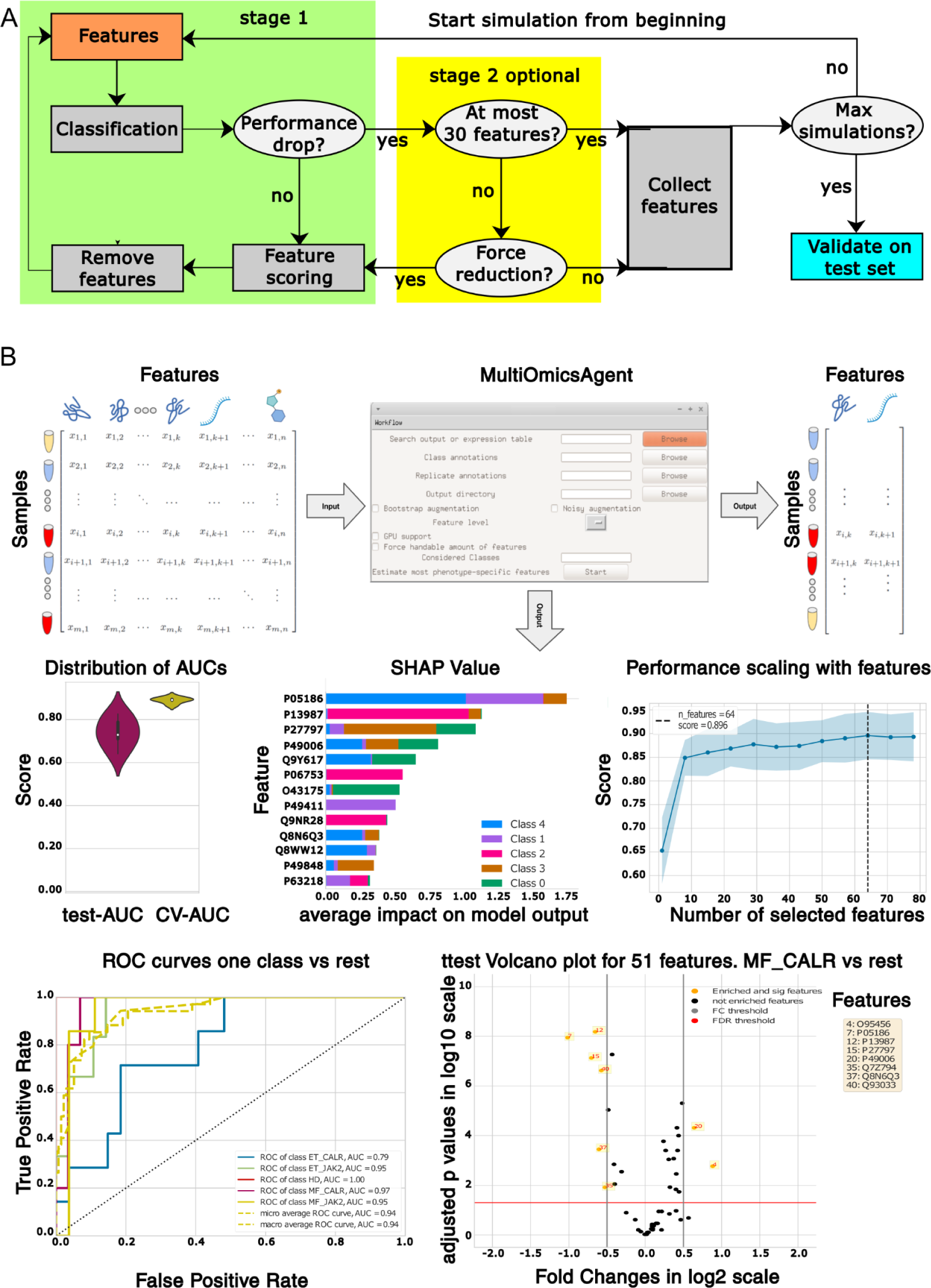
A) A flow chart illustrating how the RFE++ algorithm estimates the most phenotype-discriminating features iterating stages 1 and 2. B) The MOAgent graphical user interface (GUI) takes tabular data as input and generates an extensive output, including the most phenotype-discriminating features, model performances, and contribution evaluations of the features. Refer to the supplementary text “MOAgent supplementary visualizations’’ for a detailed description of the visualizations.

The input of MOAgent consists of:

- A csv formatted quantitative (multi-)omic matrix with samples on the rows and features (peptide/protein/transcripts/metabolites) on the columns is required. The last column specifies the classes of the samples. Note that for proteomic studies, the output of a Spectronaut [20] data analysis (exported to iq.rs report format) and Fragpipe^1^ (using a DIA_SpecLib_Quant workflow for DIA or LFQ-MBR workflow for DDA) are also directly supported as input (see supplementary figure SF1 for an example).
- A class annotation file specifying the relationship between samples and phenotypic classes. This file has to be formatted as a CSV and contains the two columns “files” and “class” (see supplementary figure SF2 for an example).
- An optional replicate annotation file specifies which samples are replicates of each other. This file has to be formatted as a CSV and contains the two columns “files” and “PatientID” (see supplementary figure SF3 for an example). When provided, this information will be used to avoid splitting related information across training, validation, and test sets, which is a very common and critical mistake.

The RFE++ algorithm is designed to find the most contributing features for phenotype classification. Since classes are often unbalanced, particular attention has been given to this aspect during the implementation of the selection procedure. For example, the area under the micro-averaged receiver operating curve (ROC) representing the true positive rate over the false positive rate for several decision thresholds (invariance of decision threshold) and classes is used to take class imbalance into account and to prevent the procedure from indiscriminately assigning samples to the most abundant class. Additionally, class distribution is maintained when generating training, validation, and test splits to achieve similar distributions across all splits. Finally, to account for the small number of samples often found in biological datasets, we also implemented a bootstrapping procedure based on repeated stratified sampling from the dataset while adding conditional Gaussian noise for each feature given the phenotype class. This allows for a proper stratified representation of all classes within each cross-validation fold used during the cross-validation training. For each sample, the bootstrapping procedure constructs two additional temporary artificial replicates within the train folds (preventing information leakage across folds due to bootstrapping which is a common issue) for each cross-validation iteration to help stabilize the algorithm.

At its core, our selection algorithm (Figure F1A) has been implemented as a Monte-Carlo-like sampling of recursive feature elimination procedures. More specifically, a randomly stratified sampled subset of the data is first set aside as a test set. The remaining data is split into a training and validation set. Subsequently, an eXtreme Gradient Boosting (XGBoost) trees classifier is trained and cross-validated on the training data using a stratified 5-fold cross-validation, and the model with the best performing features is returned. These features are ranked according to their classification contribution given by their information gain, the worst 10% are removed, and the training/validation step is repeated. In this first stage, the process of recursive feature elimination is repeated until a significant loss in classification performance is observed, which is guaranteed by the design of the algorithm. For the evaluation of the classification performance depending on the number of features, a series of scalability plots are created within each simulation for each repetition to the specified output folder and named rfecv_iter_{j}_sim_{i}_cv_roc_auc_ovo_weighted.pdf, where {j} being the rfecv repetition index and {i} the simulation iteration index. The list of features still being considered prior to the loss is saved to an output file (i.e. optimal_features_{i}.csv {i} being the simulation iteration index of stage 1) and is available to the user in the specified output folder. In a second, optional stage, if this list contains more than 30 features, the feature elimination is forced to continue till a maximum of 30 features remains. Once again, the remaining features are saved to a second output file (i.e. stage_2_forced_reduced_features_with_best_performance{i}.csv). While not necessarily optimal for classification purposes, this smaller list provides an optimal selection of features that can still be comfortably handled in biological follow-up studies. To measure the classification performance of those 30 features, a random grid hyperparameter optimized XGBoost classifier is trained in a 5-fold-cross validation mode on the train set and then validated on the validation set. The classification performance of those features can be evaluated by consulting the corresponding receiver operating curves (i.e. grid2_train_ROC_AUC_relevant_features_{i}.pdf, grid2_test_ROC_AUC_relevant_features_{i}.pdf) and confusion matrices (i.e. confusion_matrix_Grid2_train_{i}.pdf, confusion_matrix_Grid2_test_{i}.pdf) of the training and test predictions.

The entire feature selection procedure (stages 1 and 2) is repeated multiple times (three by default) after shuffling the columns of the input feature matrix and the features collected in each Monte-Carlo-like iteration.

Finally, the generalization performance is evaluated using the union of the features selected during each cycle on the set-aside test dataset after a random grid hyperparameter optimized repeated 5-fold-cross validation training of a gradient boosted decision tree classifier (i.e. final_XGB_model.json) on the initial training and validation dataset. Additionally, to provide a better overview of how the final phenotype-discriminative features (i.e. golden_features.csv) are related to the original features of the whole dataset, we also provide a csv file (i.e. significant_high_correlated_features_of_golden_features.csv) containing the Kendall correlations of those features as well as their corresponding p-values and Benjamini-Hochberg adjusted p-values of the correlations. The classification performance of the final phenotype-discriminative features is visualized in a receiver operating curve of the optimized XGBoost model (i.e. roc_auc_final_XGBclassifier_golden_features_test_performance.pdf).

In addition to the aforementioned output files, MOAgent also generates multiple graphical representations (Figure F1B) to help assess model performance and reliability.

### Setting up the MOAgent virtual machine

In line with our goal of lowering the adoption barrier of the presented workflow for users lacking coding experience, we decided to complement the pip-based package installation with a fully pre-configured virtual machine (VM) available for download as a zip-compressed file from Zenodo (https://zenodo.org/uploads/10715409). The VM seamlessly operates using the established open-source VirtualBox^2^ (version 7) software and has been tested on Intel-based macOS (>10.7), Ubuntu (versions 20.04 and 22.04), and Windows (versions 10 and 11) systems.

To use MOAgent via VM:

1. Install VirtualBox for your Operating System [https://www.virtualbox.org/].
2. Download and extract the VM file from Zenodo [https://zenodo.org/uploads/10715409].
3. Navigate to the extracted folder and double-click on the MOAgentVM.vdi file

This will start a fully prepared Xubuntu Desktop - an open-source, lightweight operating system - featuring a MOAgent desktop shortcut alongside some demo files in the “Demo” folder that can be used for testing. The user name is moagent and default password 123. To make data accessible to MOAgent, we recommend configuring the VM to either access a folder on the host machine (e.g. Windows), an external USB storage, or to use the popular pre-installed FTP client FileZilla within the VM. For more advanced users OpenSSH is also installed.

### How to use MOAgent, a demo

After the user starts the VM and logs in with the default password 123, MOAgent can be started with the desktop shortcut “MOAgent Shortcut.” The application GUI of MultiOmicsAgent will start, as shown in Figure 1B. On the Xubuntu desktop, a folder named “Demo” is located. In this folder, all the input files of the Metabolomics (GN) case study are located.

1. Use the “Browse” button of the MOAgent GUI belonging to the field “Search output or expression table” to navigate to the Demo folder on the desktop, e.g., “/home/moagent/Desktop/Demo.” Open the “input” folder and select the “metabolite_expression_table.csv” file. The Excel file content is an expression matrix with the first column, “files” containing the file names, the last column, “class” containing the class annotations, and the columns between named according to the features containing the feature expressions. An example is shown in supplementary figures SF1.
2. Use the “Browse” button of the MOAgent GUI belonging to the field “Class annotations” and navigate into the Demo folder which is located on the desktop, e.g. “/home/moagent/Desktop/Demo”. Open the “input” folder and select the “class_annotations.csv” file. An example of the class_annotations.csv file is shown in supplementary figure SF2.
3. Use the “Browse” button of the MOAgent GUI belonging to the field “Output directory” and navigate into the Demo folder, which is located on the desktop, e.g. “/home/moagent/Desktop/Demo”. Select the “output” folder.
4. Check the checkbox corresponding to “Force handable amount of features” and press the “Start” button of the MOAgent GUI to run the analysis.

The results of the MOAgent analysis will be available in the folder “/home/moagent/Desktop/Demo/output/”. They will be similar to the reported results described in GN cohort case study which can be also found in “/home/moagent/Desktop/MOAgent_supplementaries/case_studies_results/MetaboAnalyst_t ut/”.

## Results and Discussion

### MOAgent in Methylmalonyl-CoA Mutase Deficiency study: confirmations and new insights

#### (1) Transcriptome case study

We applied MOAgent to the transcriptome profile of the methylmalonyl-CoA mutase deficiency (MMA) cohort published by Forny, P. et al. [21]. In this study, 122 patients showed no, and 64 showed normal activity for the Methylmalonyl-CoA mutase (MUT) enzyme. We log2 transformed, after adding element wise the constant 1, the values in the transcriptomics expression table of the original study to achieve a more symmetrical distribution. This is generally preferable for tree-based models such as the one used by MOAgent. MOAgent achieved a test micro-averaged ROC AUC score of 76% (Figure F2A), suggesting that the list of selected transcripts was highly reliable. Among the top features in the SHAP summary plot, we found the MUT transcript (ENSG00000146085) (Figure F2B), also identified as a key disease-relevant transcript in the original study.

**Figure F2:**
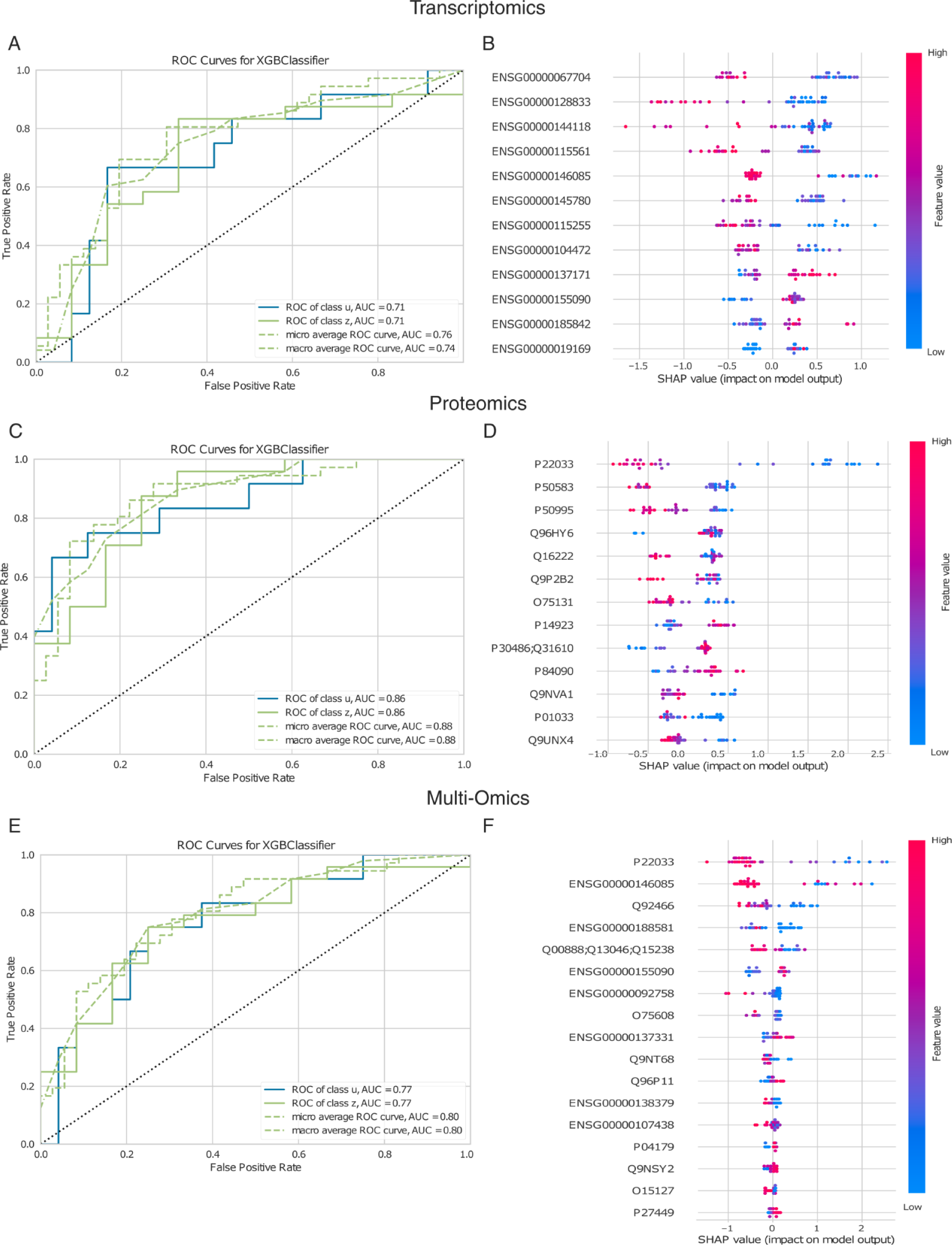
A, C, D show respectively the ROC curves for the test set of the MUT active and inactive classification task based on transcript, protein and transcript combined with protein (multi omic) expressions. The corresponding micro average ROC curve AUC scores are 76%, 88% and 80%, respectively and indicate across all omics levels a high reliability of the extracted most classification-relevant features from the dataset. B, C, F show the corresponding SHAP summary plots and visualize the classification contribution of the extracted features, where features with a higher SHAP value deviation indicate a stronger class discriminatory power.

#### (2) Protein case study

We further applied a similar workflow to the proteomics data published in the same study by Forny et al. The test micro averaged ROC AUC score for the classification was 88% (Figure F2C). Also, in this modality, the protein corresponding to the MUT gene (UniProt P22033) is selected and ranked first (Figure F2D).

#### (3) Multi-Omics case study

Finally, we applied MOAgent on the concatenated transcriptomics and proteomics data from the MMA study. The test micro averaged ROC AUC score of the classification was 80% (Figure F2E). Once again, P22033 and ENSG00000146085 were among the top-selected features (Figure F2F).

### MOAgent easily retrieves known biomarker-candidates of a complex myeloproliferative neoplasms (MPN) cohort

We applied MOAgent on the myeloproliferative neoplasms (MPN) cohort published by Wildschut et al. [19]. This study includes multiple phenotypic classes of blood cancer disease. The two major forms of MPN (Myelofibrosis (MF) and essential thrombocythemia (ET)), are mainly driven by mutations in the CALR and the JAK2 genes. Additionally, blood from healthy donors (HD) with matching gender and age were used as a control class. The five phenotypic classes were, therefore, designated HD, ET CALR, ET JAK2, MF CALR, and MF JAK2, respectively.

In the original publication, we identified 21 proteins that could be used to achieve high classification accuracy between the five phenotypes. Here, we first applied the DIA_SpecLib_Quant^3^ workflow of Fragpipe - a Java GUI that provides a comprehensive suite of computational tools for analyzing mass spectrometry-based proteomics data - on the raw MS files from the study (dataset PXD036075 on ProteomeXchange) to generate a file capturing the protein expressions (i.e. diann.tsv). In combination with the patient- and class-annotation CSV files to derive, with one click, a list of phenotype-specific proteins via MOAgent (see supplementary Table 1). Three (i.e. NEK5, STYX, and ZNF735) of the 21 proteins selected in the original study were not identified by Fragpipe. The output of MOAgent contained 15 highly correlated or identical proteins for the remaining 18. Notably, while the original analysis was performed with Spectronaut - a commercial software package developed for analyzing Data-Independent Acquisition (DIA) proteomics experiments - and included an ad-hoc programmed batch regression, missing value imputation, and feature selection pipeline, our reanalysis entirely relied on open-source software. It completely bypassed the need to write code for the analysis. It should also be mentioned that in the original publication, we used an earlier implementation of the RFE that optimized Extreme Randomized Decision Trees (Extra Trees) towards the macro averaged F1 score (i.e. the class averaged harmonic mean between precision and recall at one specific decision threshold), whereas MOAgent uses an XGBoost classifier and optimizes the model regarding micro averaged ROC AUC, considering the trade-off of true positives over false positives for a range of decision thresholds. Therefore, the newer implementation provides a classification threshold-independent evaluation and, thus a more meaningful assessment of the selected features’ discriminatory power. All in all, 83% of the class-relevant proteins detected in the original study were identified or highly kendall correlated alternatives (after multi testing correction with Benjamini-Hochberg correction) of those by MOAgent with a test micro averaged ROC AUC score of 94% (Figure F3A, F3B) as highly class-relevant without needing to write a single line of code.

**Figure F3:**
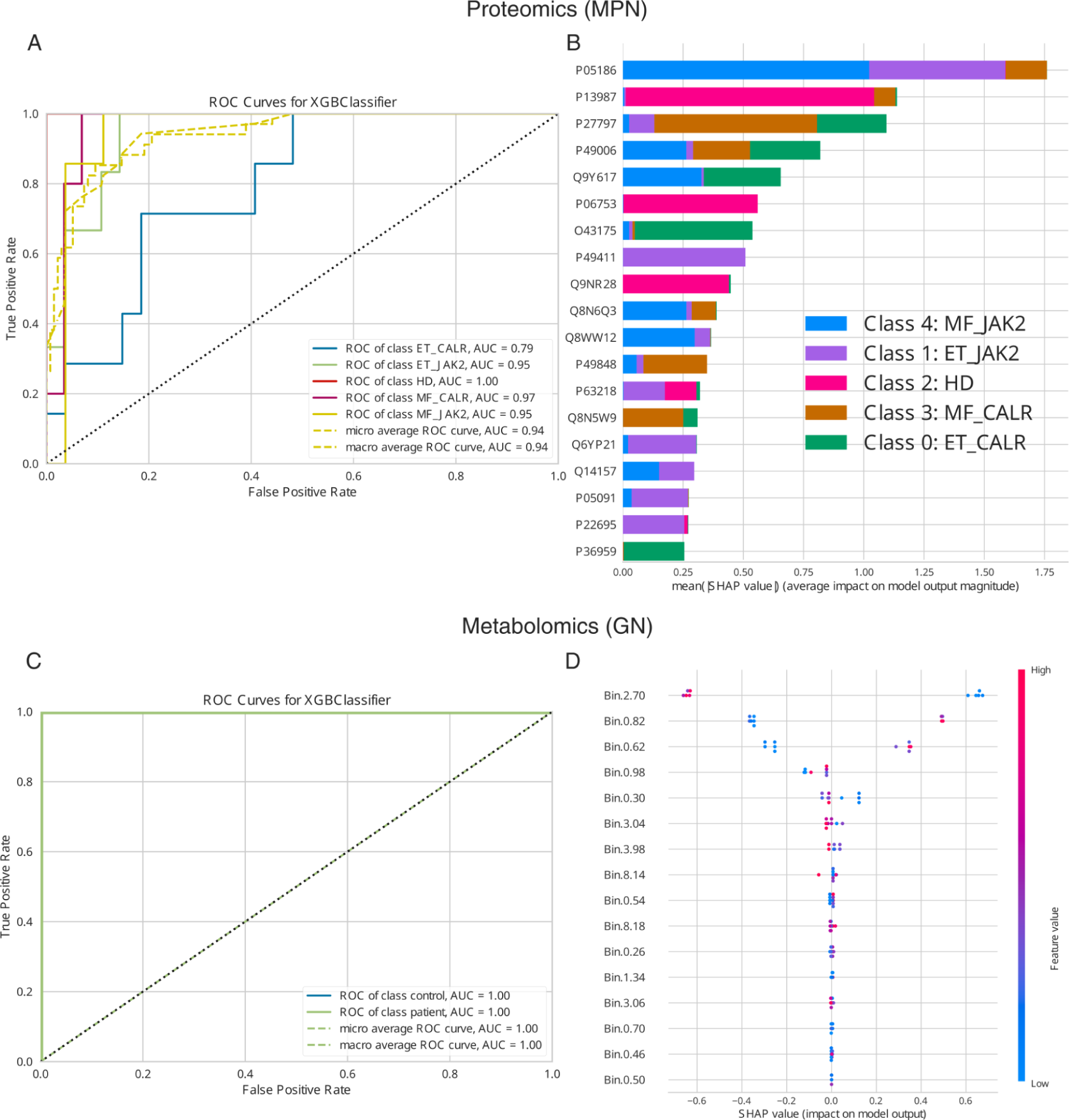
A, shows the ROC curves for the test set of the myeloproliferative neoplasms phenotypic classification task based on protein expressions. The corresponding micro average ROC curve AUC scores 94% and indicates a high reliability of the extracted most phenotype classification-relevant features from the dataset. B, shows the corresponding SHAP summary plot and visualizes the classification contribution of the extracted features of the MPN classification task. C, shows the ROC curves for the test set of the Glomerulonephritis classification task based on metabolite expressions. The corresponding micro average ROC curve AUC scores 100% and indicates a high reliability of the extracted most classification-relevant features from the dataset. D, shows the corresponding SHAP summary plot and visualizes the classification contribution of the extracted features of the GN classification task.

### MOAgent perfectly classified Glomerulonephritis using NMR data, pinpointing citrate as crucial, without user coding

We applied MOAgent to a metabolite expression table generated by Nuclear Magnetic Resonance (NRM) for the classification problem that is provided on the MetaboAnalyst [11] website. The classification tutorial from this website contains 25 healthy controls and 25 Glomerulonephritis (GN) affected patients, a subset from the study of Psihogios et al. [22]. The super bin 2.70 ppm corresponding to citrate is known from the study as a key player in the disease and was identified as top feature in the MOAgent SHAP summary plot (Figure F3C). For this study, the test micro averaged ROC AUC score of the classification was 100% (Figure F3D). Once again, these results were achieved without writing a single line of code by the user.

### MOAgent validated cell clonality in multiple myeloma with high accuracy, highlighting Immunoglobulin light chains as key features

We applied MOAgent to validate the clonality of CD138-sorted cells from a multiple myeloma patient cohort published by Kropivsek, K. et al. [23]. The quantitative CD138 protein data matrix contained a total of 74 samples and two phenotypic classes (i.e. 48 samples with Kappa light-chain and 26 with Lambda light-chain). MOAgent achieved test micro averaged ROC AUC scores of 91% on the protein level (Figure F4A) and 95% on the peptide level (Figure F4C). As expected, at the protein level in the SHAP summary plot, we find features of the Immunoglobulines Kappa and Lambda light chain as the two most significant hits (Figure F4B). Similarly at the peptide level, we find peptides from the Immunoglobulines Kappa and Lambda among the top-ranking features for classification (Figure F4D). Once again, these results were achieved without a single line of code.

**Figure F4:**
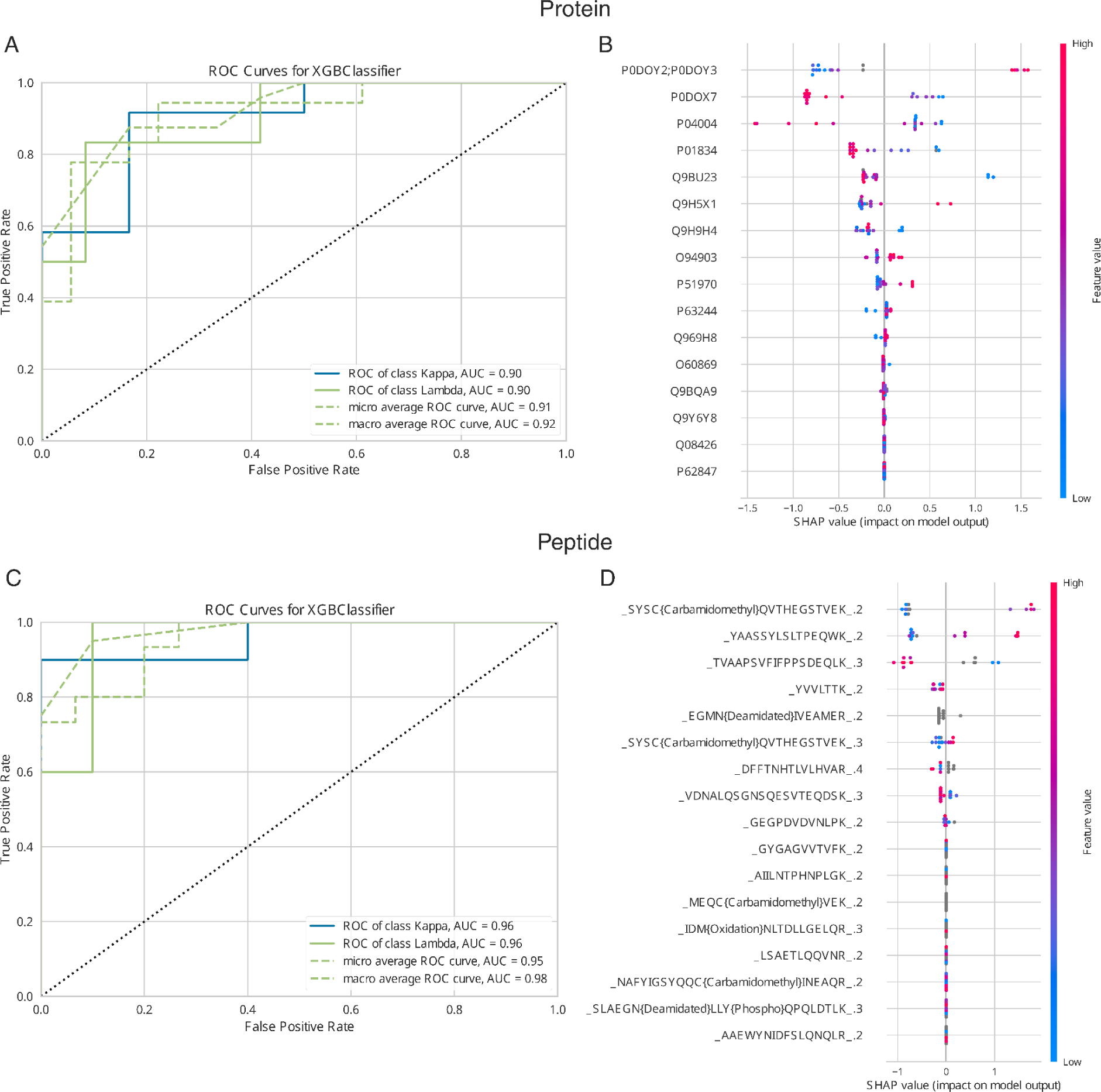
A and C, show the ROC curves for the test set of the clonality in multiple myeloma (Kappa-Lambda) classification task based on protein and peptide expressions, respectively. The corresponding micro average ROC curve AUC scores are 91% and 95%, indicating a high reliability of the extracted most classification-relevant features from the dataset. B and D, show the corresponding SHAP summary plots and visualize the classification contribution of the extracted features of the Kappa-Lambda classification task.

### New insights from the case studies analysis using MOAgent

The integration of MOAgent within the realms of protein and transcriptomic analysis, or broadly across multiple omic levels, has unveiled novel perspectives on the comparative contributions of various omic features to phenotypic manifestations. In studies focusing on proteins and transcriptomes, the AUC test scores revealed a nuanced balance between true positive and false positive allocations of samples to phenotypic categories, with protein expressions yielding a superior trade-off at 88%, compared to 76% in the transcriptomic analyses of the MMA cohort. In the Kappa-Lambda case study an improvement of 4% on the peptide level compared to protein level can be observed strengthen the perspective to focus on peptide level for downstream proteomics analysis as already discussed in Plubell et al. [24].

MOAgent allows not only multi-omics studies, it enables also multi-class studies in the same framework as showcased in the MPN cohort example. Figure F3A reveals the ET CALR class is more challenging to classify than any other of the classes, which can be concluded by the class specific ROC curves MOAgent provides.

When examining multi-omic studies, MOAgent facilitates a unique confrontation among features from disparate omic strata, highlighting the most discriminative attributes for class separation. Notably, the MMUT corresponding protein in the MMA study and its transcript emerge as leading indicators, as determined by SHAP values. This cross-omic rivalry uncovers biologically significant features that might elude detection within isolated omic investigations. For instance, a transcript deemed critical for classification may not translate into a discernible protein expression within phenotypic variations, despite its association with a protein complex that includes a more influential protein member. Such discrepancies might stem from post-translational modifications or insufficient expression variances across phenotypic classes, given the sample size. Therefore, MOAgent’s SHAP-informed feature ranking presents an innovative lens for dissecting phenotypic nuances on a systemic biological scale supporting advances towards biological digital twins through multi-omic integration.

Further insights from the MMA cohort underscore the strategic value under certain circumstances of prioritizing protein expressions, which achieved an AUC score of 88%, surpassing the 80% score from a combined protein-transcriptomic approach.

This finding suggests that, depending on the objectives of a study, significant resources could be conserved by focusing on a single omic dimension, thereby streamlining the analytical process without necessarily compromising the depth of phenotypic understanding. Nevertheless, the incremental improvement observed in the MMA study, where the integration of protein and transcript expressions enhanced the true positive versus false positive ratio by 4%, underscores the potential of multi-omic approaches to refine phenotypic classifications beyond the capabilities of single omic analyses.

### Automatic provided data and result visualizations by MOAgent

#### (1) Feature Analysis and Visualization

(Figure SF4 and SF5) MOAgent generates both UMAP (Uniform Manifold Approximation and Projection) (see supplementary Figures SF4) and PCA (Principal Component Analysis) (see supplementary Figures SF5) plots, enabling a conclusion towards non-linear and linear separability of data points regarding phenotypic classes, respectively. Conclusions considering similarities between different clusters should be taken from PCA plots, since the orthogonal mapping and projection onto the most variance contributing components (principle components) are more euclidean distance preserving than the two UMAP components.

These plots are created considering the entire spectrum of features as well as focusing exclusively on the most phenotype-discriminating features and allows a qualitative investigation of class distangeling after the feature reduction. For successful feature selection, we anticipate the PCA and UMAP plots, when focused on phenotype-discriminating features, to exhibit less mixing and more structured clustering aligned with the phenotype classes of the dataset.

#### (2) Performance and Reliability Evaluation

(Figure F5) To assess the classification performance and its generalization ability, MOAgent creates violin plots incorporating box plots. These plots visualize the distribution, quantiles, and medians of ROC (Receiver Operating Characteristic) and AUC (Area Under the Curve) scores for both cross-validation and test sets throughout the feature selection process (stages 1 and 2). Additionally, test scores for accuracy (ACC), macro averaged F1 score (MAVG_F1), which corresponds to averaging the unweighted mean per label, and weighted average F1 score (WAVG_F1), which corresponds to averaging the support-weighted mean per label, are visualized. Note that the decision thresholds of the classifier are optimized on the cross-validation set for maximum sensitivity (true positive rate) while minimizing the false positive rate. This decision threshold represents the best trade-off between correctly identifying positive cases while avoiding false positives. It corresponds to the point on the ROC curve that has the shortest distance to the top left corner (0,1) in the cartesian coordinate system. However, we strongly recommend to focus on cross-validation and test AUC distributions to evaluate the features phenotype prediction capabilities.

**Figure F5:**
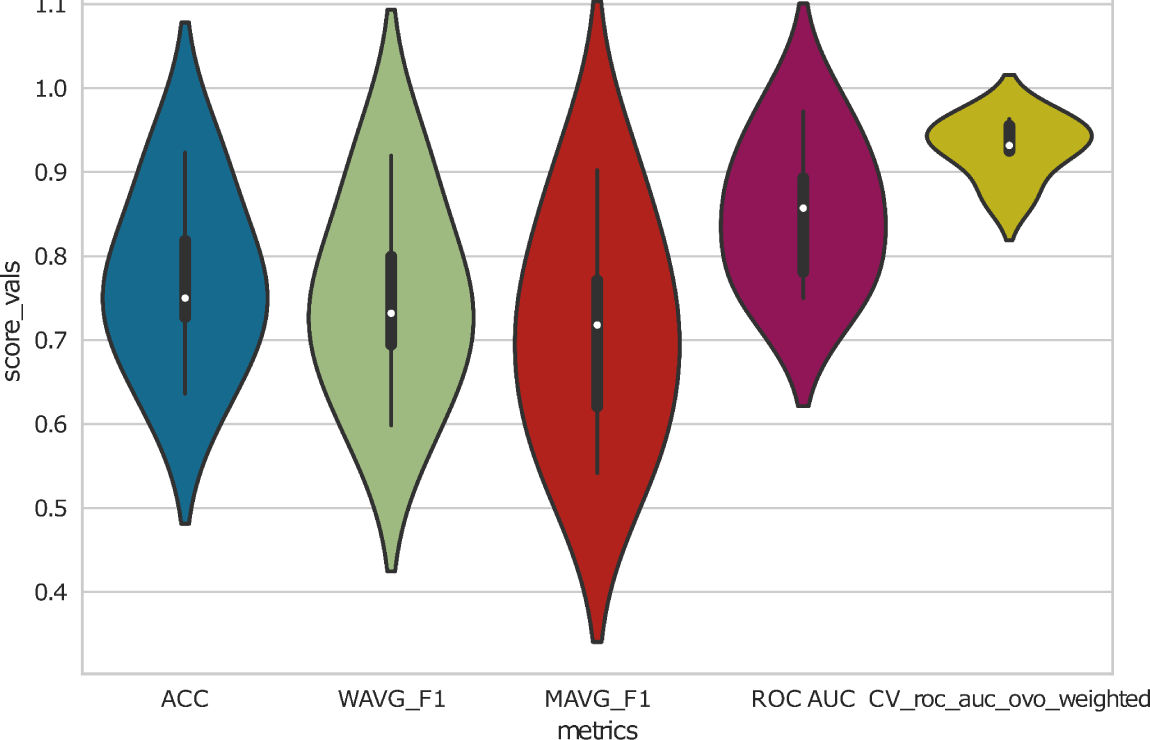
In each Monte-Carlo-likesimulation, a random grid hyperparameter optimized XGBoost classifier (named Grid2) is trained in a 5-fold cross-validation mode on a train set and additionally tested on a separate test set. This figure from the Kappa-Lambda case study is shown as an example and visualizes the evaluation scores’ distributions in violin plots and box plots.

#### (3) ROC Curve Analysis

(Figure F6 A and B) ROC curves are provided, illustrating the true positive rate against the false positive rate at various decision thresholds. This analysis is conducted for the XGBoost classifier in the final model and for training (Figure F6A) and test set (Figure F6B). The area under these curves serves as an indicator of model performance and should be preferred over confusion matrices to evaluate the predictive power of selected features. The bigger the area under the ROC curve, the better the model performs and the more reliable the selected features are.

**Figure F6:**
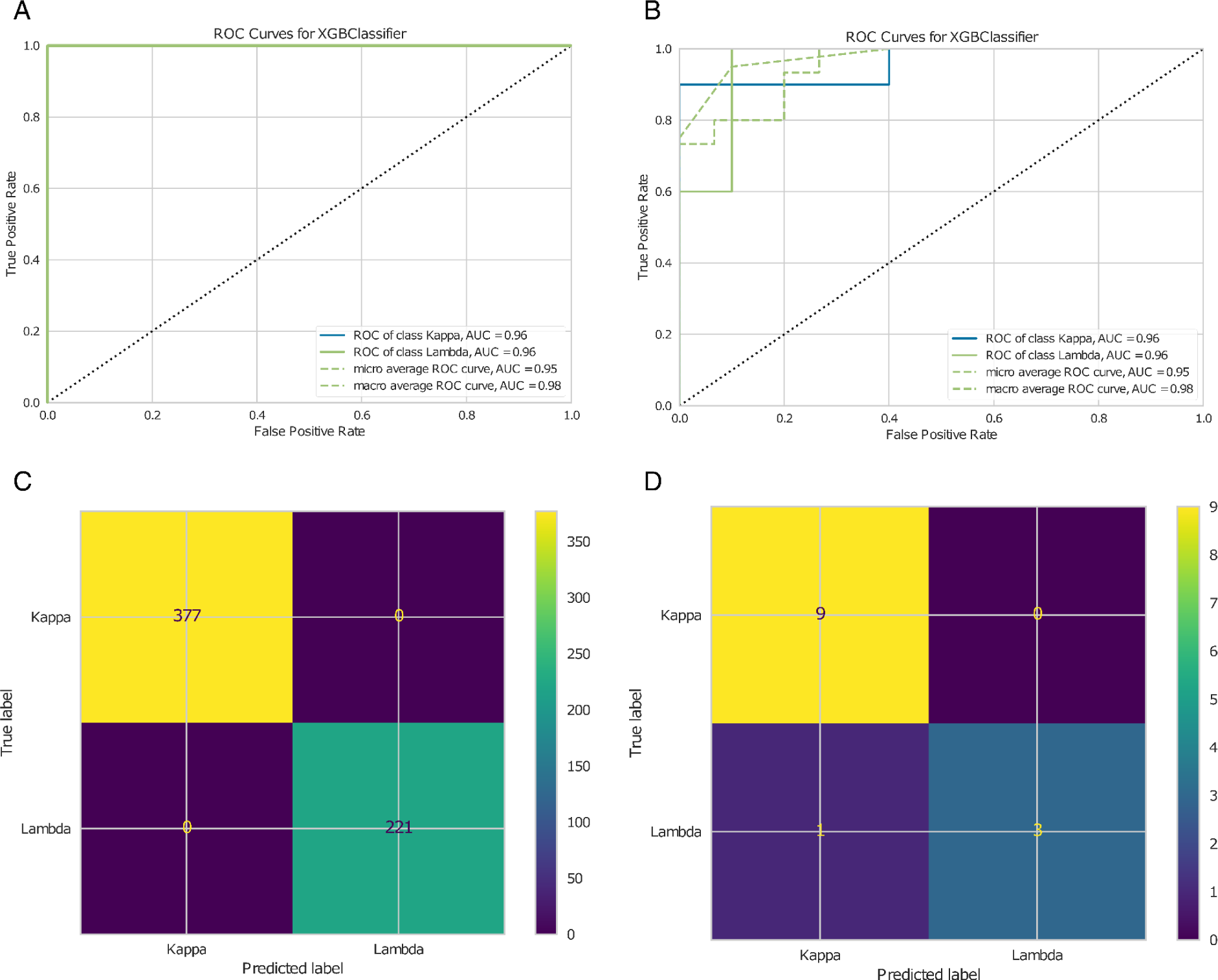
A and B, show example ROC curve for training and test predictions of the Kappa-Lambda peptide case study, respectively. C and D, show example figures for the train and test confusion matrices of the Kappa-Lambda peptide case study.

The roc_auc_final_XGBclassifier_golden_features_test_performance.pdf is the first of two key visual outputs a user should pay special attention to. Together with xgb_final_features_shap_test_performance.pdf (see “feature trustability visualization”), it illustrates the reliability of the selected features.

#### (4) Confusion Matrix and Further Statistics

(Figure F6 C and D) Confusion matrices are generated in each simulation for train (Figure F6C) and test (Figure F6D) predictions of the hyperparameter tuned XGBmodel, allowing for the computation of additional statistical measures besides the ones provided, like accuracy, F1 score, sensitivity, and specificity, which are pertinent to the specific application. The better the classifier performs, the closer this matrix is to a diagonal matrix (all off-diagonal entries tend to be zero). The provided confusion matrices correspond to the final model using the decision threshold for which the train and test ROC curves respectively were closest to the top left corner (0,1) in the cartesian system representing the true positive rate over the false positive rate.

#### (5) Feature Trustability Visualization

(supplementary Figure SF6) Visualizations for direct measurement of the selected features’ reliability are provided through classical volcano plots (see supplementary SF6A) (after a data-dependent minimum value imputation for missing values) using two-sided Mann-Whitney U tests for each feature and SHAP (SHapley Additive exPlanations) (see supplementary SF6B) value plots. The volcano plots are computed for all features and specifically for selected features, with the latter being less stringent towards multi-testing correction. Notably, both utilize Benjamini-Hochberg-adjusted p-values. In a multiclass phenotype classification scenario, multiple volcano plots (e.g., five for a five phenotype-class problem) are generated, each comparing one phenotypic class against the rest. If a user specifies only two classes in the GUI to investigate, then rest corresponds to a single comparison class. The SHAP value plot visualizes how much a feature contributed to the classification task, with more relevant features having bigger absolute values. The file xgb_final_features_shap_test_performance.pdf is the second of the two key visual outputs and shows the classification importance of the features in golden_features.csv.

#### (6) Correlation Analysis and Feature Distribution

(supplementary Figure SF7, SF8 and SF9) The Kendall correlations of selected features are visualized in a heatmap (see supplementary SF7). The Kendall correlation is chosen, because it does not assume a specific feature distribution as a person correlation does for example. Additionally, non-linear relationships can be reliably captured. Kendall is also suitable for smaller datasets and is less sensitive to ties or very close feature values. A hierarchical clustering with heatmap across the selected features is provided (see supplementary SF8). MOAgent additionally provides boxplots (see supplementary SF9) that visualize the distribution of feature expression values across different classes, as well as a heatmap showing the dataset’s expression patterns for the most phenotype-discriminating features.

#### (7) Feature classification performance scalability

(Figure F7) In stage 1, an iteration of recursive feature eliminations is applied, and in each simulation for each recursive feature elimination repetition, a scalability plot is generated. One scalability plot shows the mean ROC AUC classification performance with standard deviation over the number of features.

**Figure F7:**
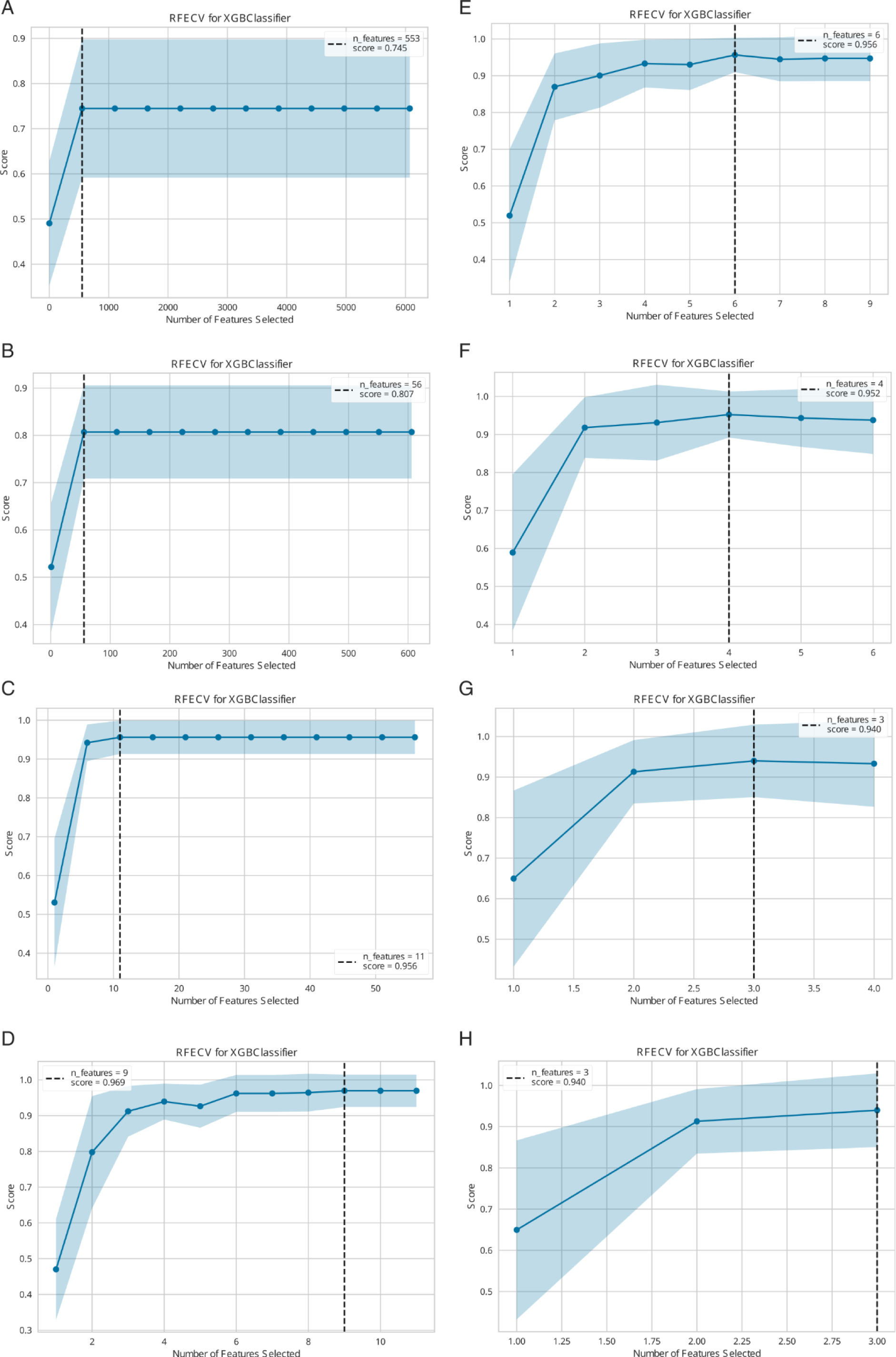
Scalability plots for the Kappa-Lambda case study on protein level. The iterations are sorted from top to down and left to right (A-H), indicated by the reducing maximum number of considered features from plot to plot.

## Conclusion

While it has never been easier to identify and quantify thousands of molecular entities such as transcripts, proteins, and metabolites from biological samples, the extraction of biological significance from these multimodal datasets remains challenging and mostly confined to the realm of expert data scientists. Among the challenges that need to be addressed are the necessity to: handle missing values; consider complex combinations of multiple molecular makeups (biomarker signatures); and efficiently control the false discovery rate (FDR) in the presence of limited replicates and large feature numbers. Machine Learning offers promising approaches to tackle this task if (as a rule of thumb) at least twenty data points per class in the dataset are available. However, the accessibility of these methodologies is often restricted by high requirements in coding skills and ML domain knowledge. MOAgent addresses these concerns by providing access to an advanced, ML-based, multi-omic biomarker candidate discovery pipeline via a simple, cross-platform supported, user-centric GUI. Overall, by simplifying and optimizing the biomarker candidate discovery process, MOAgent promises a harmonious blend of accuracy and accessibility, which will push the boundaries of interdisciplinary omic research.

## Author contributions

Jens Settelmeier conducted the project, wrote the first draft of the manuscript, developed and implemented the algorithms in MOBiceps, including RFE++, MOAgent, created the figures, performed the re-analysis of the case study, including the search with Fragpipe and post-analysis with MOAgent, set up the GitHub repositories, and is the maintainer of MOAgent and MOBiceps. Sandra Goetze tested and provided feedback regarding the usability of MOAgent and the results of the RFE++. Sandra also reviewed the manuscript. Julia Boshart generated data that was crucial for testing. Julia measured samples which were used to test MOAgent, reviewed the manuscript and contributed to the discussions from a biology perspective. Jianbo Fu contributed with manuscript revisions and technical discussions regarding biomarker-candidate discovery. Sebastian Steiner reviewed the manuscript and was involved in discussions, contributing with biology expertise. Martin Gesell reviewed the manuscript and was involved in discussions, providing biology expertise. Diyora Salimova contributed detailed feedback regarding the algorithmic design, which involved machine learning and statistics discussions. Diyora also reviewed the manuscript. Peter Schüffler reviewed the manuscript and contributed to the machine-learning discussions. Patrick Pedrioli co-supervised the project and contributed with code and manuscript revisions, and figure discussions. Further, Patrick contributed heavily to the algorithmic design discussions. Bernd Wollscheid and Patrick Pedrioli co-supervised the project, acquired funding, and supported the development of manuscript with critical feedback at all stages.

## Acknowledgements

J.S. acknowledges fruitful discussions with members of the Wollscheid lab and the ETH AI Center. Special thanks go to Jacqueline Hammer for testing the installation guide and to Ruedi Aebersold for helpful discussions.

## Funding

This work was supported by funds from the strategic focus area Personalized Health and Related Technologies from the ETH Domain [2022/601] to S.G. and P.G.A.P.

## Conflict of Interest Statement

The authors declare no competing financial interests.

## Footnotes

1 https://github.com/Nesvilab/FragPipe

2 https://www.virtualbox.org/

3 >https://fragpipe.nesvilab.org/docs/tutorial_DIA.html

## Notes

https://zenodo.org/uploads/10715409

https://github.com/Wollscheidlab/MOAgent

https://github.com/Wollscheidlab/MOBiceps

### Competing Interest Statement

The authors have declared no competing interest.

## References

1. Osipov A, Nikolic O, Gertych A, et al. The Molecular Twin artificial-intelligence platform integrates multi-omic data to predict outcomes for pancreatic adenocarcinoma patients. Nat. Cancer 2024; 5:299–314

2. Aebersold R, Mann M. Mass spectrometry-based proteomics. Nature 2003; 422:198–207

3. Ong S-E, Mann M. Mass spectrometry–based proteomics turns quantitative. Nat. Chem. Biol. 2005; 1:252–262

4. Bantscheff M, Schirle M, Sweetman G, et al. Quantitative mass spectrometry in proteomics: a critical review. Anal. Bioanal. Chem. 2007; 389:1017–1031

5. Rosenberger, G. et al. (2017). Inference and quantification of peptidoforms in large sample cohorts by swath-ms. Nature biotechnology, 35(8), 781– 788

6. Xuan Y, Bateman NW, Gallien S, et al. Standardization and harmonization of distributed multi-center proteotype analysis supporting precision medicine studies. Nat. Commun. 2020; 11:5248

7. Rozanova S, Barkovits K, Nikolov M, et al. Quantitative Mass Spectrometry-Based Proteomics: An Overview. Methods Mol. Biol. Clifton NJ 2021; 2228:85–116

8. Cherkaoui S, Durot S, Bradley J, et al. A functional analysis of 180 cancer cell lines reveals conserved intrinsic metabolic programs. Mol. Syst. Biol. 2022; 18:e11033

9. Babu M, Snyder M. Multi-Omics Profiling for Health. Mol. Cell. Proteomics MCP 2023; 22:100561

10. DeGroat W, Mendhe D, Bhusari A, et al. IntelliGenes: a novel machine learning pipeline for biomarker discovery and predictive analysis using multi-genomic profiles. Bioinformatics 2023; 39:btad755

11. Xia, J. and Wishart, D. S. (2016). Using metaboanalyst 3.0 for comprehensive metabolomics data analysis. Current protocols in bioinformatics, 55(1), 14–10

12. López-Fernández, H. et al. (2015). Mass-up: an all-in-one open software application for maldi-tof mass spectrometry knowledge discovery. BMC bioinformatics, 16, 1–12

13. Argelaguet R, Velten B, Arnol D, et al. Multi-Omics Factor Analysis-a framework for unsupervised integration of multi-omics data sets. Mol. Syst. Biol. 2018; 14:e8124

14. Shi Z, Wen B, Gao Q, et al. Feature Selection Methods for Protein Biomarker Discovery from Proteomics or Multiomics Data. Mol. Cell. Proteomics 2021; 20:100083

15. Benjamin KJM, Katipalli T, Paquola ACM. dRFEtools: dynamic recursive feature elimination for omics. Bioinforma. Oxf. Engl. 2023; 39:btad513

16. Smit S, Hoefsloot HCJ, Smilde AK. Statistical data processing in clinical proteomics. J. Chromatogr. B Analyt. Technol. Biomed. Life. Sci. 2008; 866:77–88

17. Guyon I, Weston J, Barnhill S, et al. Gene Selection for Cancer Classification using Support Vector Machines. Mach. Learn. 2002; 46:389–422

18. Goetze S, Frey K, Rohrer L, et al. Reproducible Determination of High-Density Lipoprotein Proteotypes. J. Proteome Res. 2021; 20:4974–4984

19. Wildschut, M. H. et al. (2023). Proteogenetic drug response profiling elucidates targetable vulnerabilities of myelofibrosis. Nature Communications, 14(1), 6414

20. Bernhardt, O. M. et al. (2012). Spectronaut: a fast and efficient algorithm for mrm-like processing of data independent acquisition (SWATH-MS) data

21. Forny P, Bonilla X, Lamparter D, et al. Integrated multi-omics reveals anaplerotic rewiring in methylmalonyl-CoA mutase deficiency. Nat. Metab. 2023; 5:80–95

22. Psihogios NG, Kalaitzidis RG, Dimou S, et al. Evaluation of tubulointerstitial lesions’ severity in patients with glomerulonephritides: an NMR-based metabonomic study. J. Proteome Res. 2007; 6:3760–3770

23. Kropivsek K, Kachel P, Goetze S, et al. Ex vivo drug response heterogeneity reveals personalized therapeutic strategies for patients with multiple myeloma. Nat. Cancer 2023; 4:734–753

24. Plubell DL, Käll L, Webb-Robertson B-J, et al. Putting Humpty Dumpty Back Together Again: What Does Protein Quantification Mean in Bottom-Up Proteomics? J. Proteome Res. 2022; 21:891–898

