## Supplementary Material for "MultiOmicsAgent: Guided extreme gradient-boosted decision trees-based approaches for biomarker-candidate discovery in multi-omics data"

1) Supplementary Figures

    Figures: SF1 - SF9

2) Supplementary Tables

    Table 1

### 1. Supplementary Figures

#### Example input files

metabolite_expression_table.csv

| | A | B | C | D | E | |
|---|---|---|---|---|---|---|
| 1 | files | Bin.9.98 | Bin.9.94 | Bin.9.90 | Bin.9.86 | Bi |
| 2 | P002 | 2.00E-05 | 2.00E-05 | 2.00E-05 | 2.00E-05 | |
| 3 | P012 | -3.00E-05 | -3.00E-05 | -3.00E-05 | -1.00E-05 | |
| 4 | P014 | 0.00024 | 0.00017 | 0.00016 | 0.00018 | |
| 5 | P027 | 9.00E-05 | 0.00013 | 9.00E-05 | 9.00E-05 | |
| 6 | P034 | 0.00012 | 0.00015 | 2.00E-05 | 3.00E-05 | |
| 7 | P037 | 1.00E-05 | -1.00E-05 | 2.00E-05 | -3.00E-05 | |
| 8 | P038 | 1.00E-05 | 1.00E-05 | 1.00E-05 | -5.00E-05 | |
| 9 | P041 | -9.00E-05 | -6.00E-05 | -7.00E-05 | 1.00E-05 | |
| 10 | P042 | 0.00015 | 8.00E-05 | 0.00017 | 3.00E-05 | |

**Figure SF1:** Expression table, with the first column "files" corresponding to the sample file identifiers, the last column "class" corresponding to the sample class assignment, and all other columns named by the features corresponding to the feature expressions.

class_annotations.csv

| | A | B |
|---|---|---|
| 1 | files | class |
| 2 | P002 | patient |
| 3 | P012 | patient |
| 4 | P014 | patient |
| 5 | P027 | patient |
| 6 | P034 | patient |
| 7 | P037 | patient |
| 8 | P038 | patient |
| 9 | P041 | patient |
| 10 | P042 | patient |

**Figure SF2:** Class annotation table, with the first column "files" corresponding to the samples' file identifiers and the column "class" corresponding to their class assignments.

| | A | B |
|---|---|---|
| 1 | files | PatientID |
| 2 | wildschm_B1909_024.mzML | 14 |
| 3 | wildschm_B1909_025.mzML | 72 |
| 4 | wildschm_B1909_026.mzML | 100 |
| 5 | wildschm_B1909_027.mzML | 55 |
| 6 | wildschm_B1909_029.mzML | 7 |
| 7 | wildschm_B1909_030.mzML | 98 |
| 8 | wildschm_B1909_032.mzML | 15 |
| 9 | wildschm_B1909_033.mzML | 17 |
| 10 | wildschm_B1909_036.mzML | 108 |
| 11 | wildschm_B1909_037.mzML | 58 |
| 12 | wildschm_B1909_039.mzML | 94 |
| 13 | wildschm_B1909_040.mzML | 5 |
| 14 | wildschm_B1909_042.mzML | 76 |
| 15 | wildschm_B1909_043.mzML | 82 |
| 16 | wildschm_B1909_044.mzML | 81 |
| 17 | wildschm_B1909_045.mzML | 57 |
| 18 | wildschm_B1909_046.mzML | 104 |
| 19 | wildschm_B1909_049.mzML | 85 |
| 20 | wildschm_B1909_050.mzML | 99 |

**Figure SF3:** (Optional) patient annotation table with first column "files" corresponding to the file identifiers of the samples and the column "PatientID" to the individual the sample belongs to. This tabular file is used if several files (replicates) belong to the same individual (e.g. patient). Several files can be assigned the same PatientID, to let the algorithm know, the files are replicates of the same patient (individual).

### Feature Analysis and Visualization

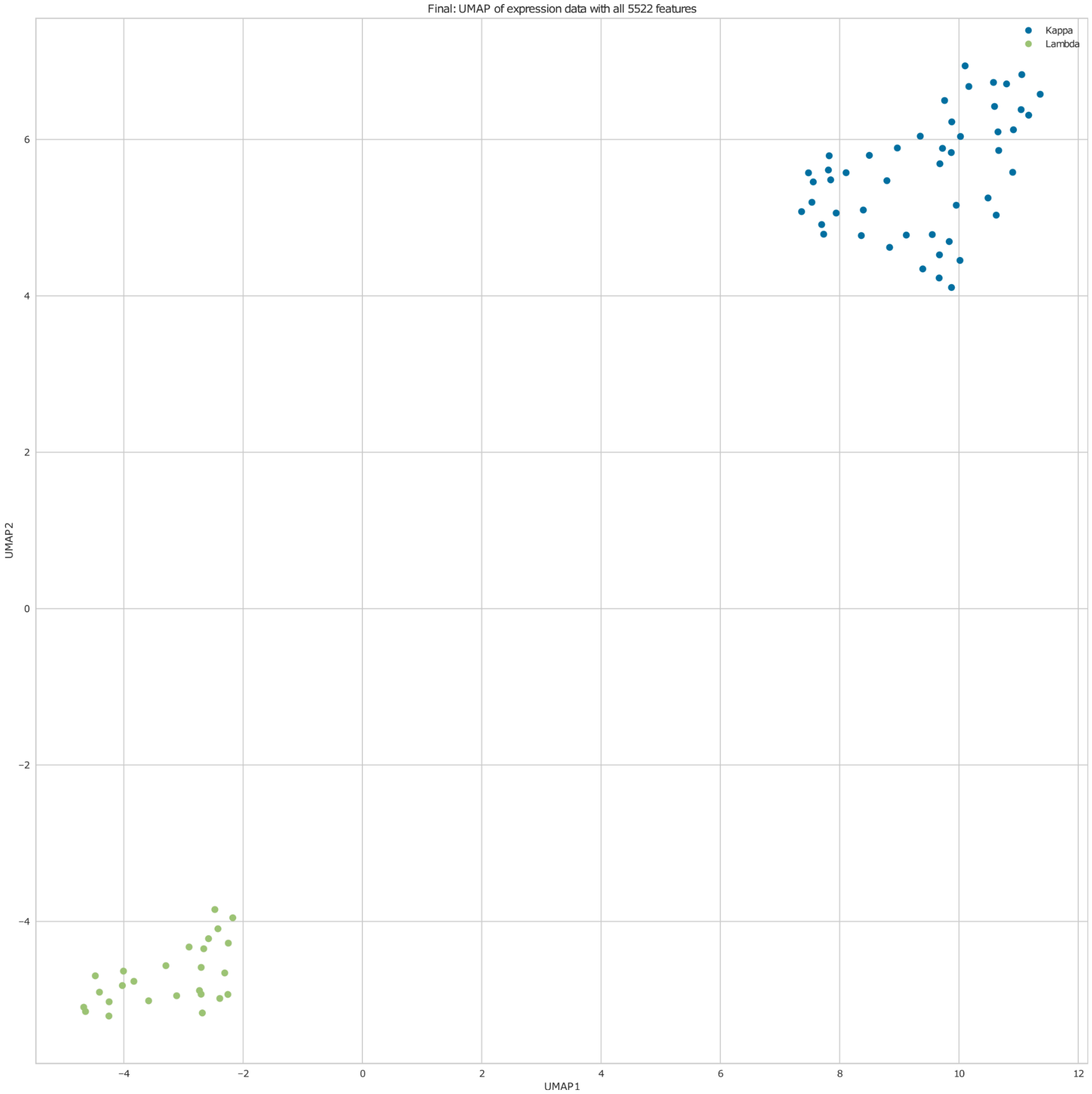


**Figure SF4:** UMAP using all features of the Kappa-Lambda Protein case study.

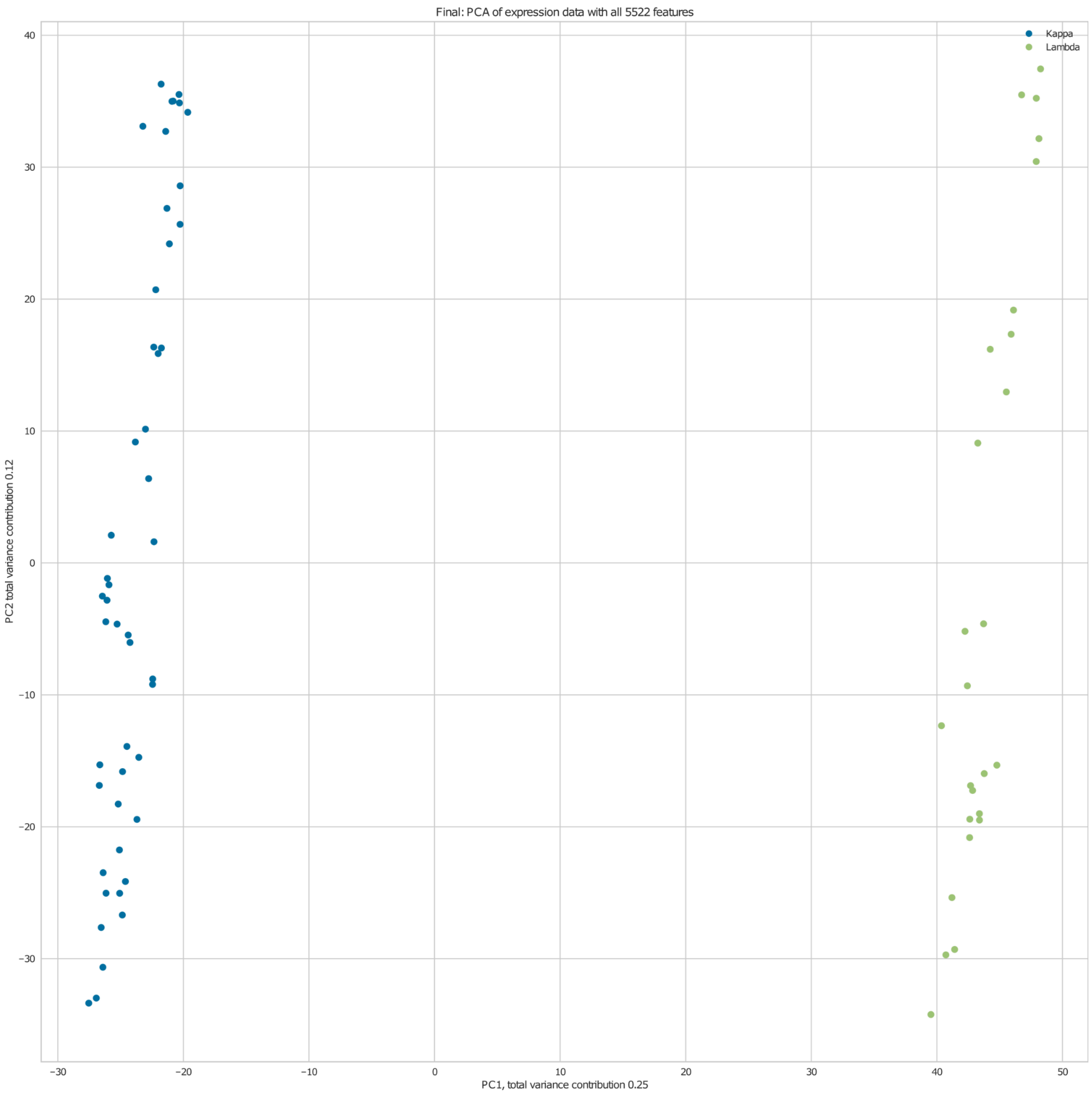


**Figure SF5:** PCA using all features of the Kappa-Lambda Protein case study.

### Feature Trustability Visualization

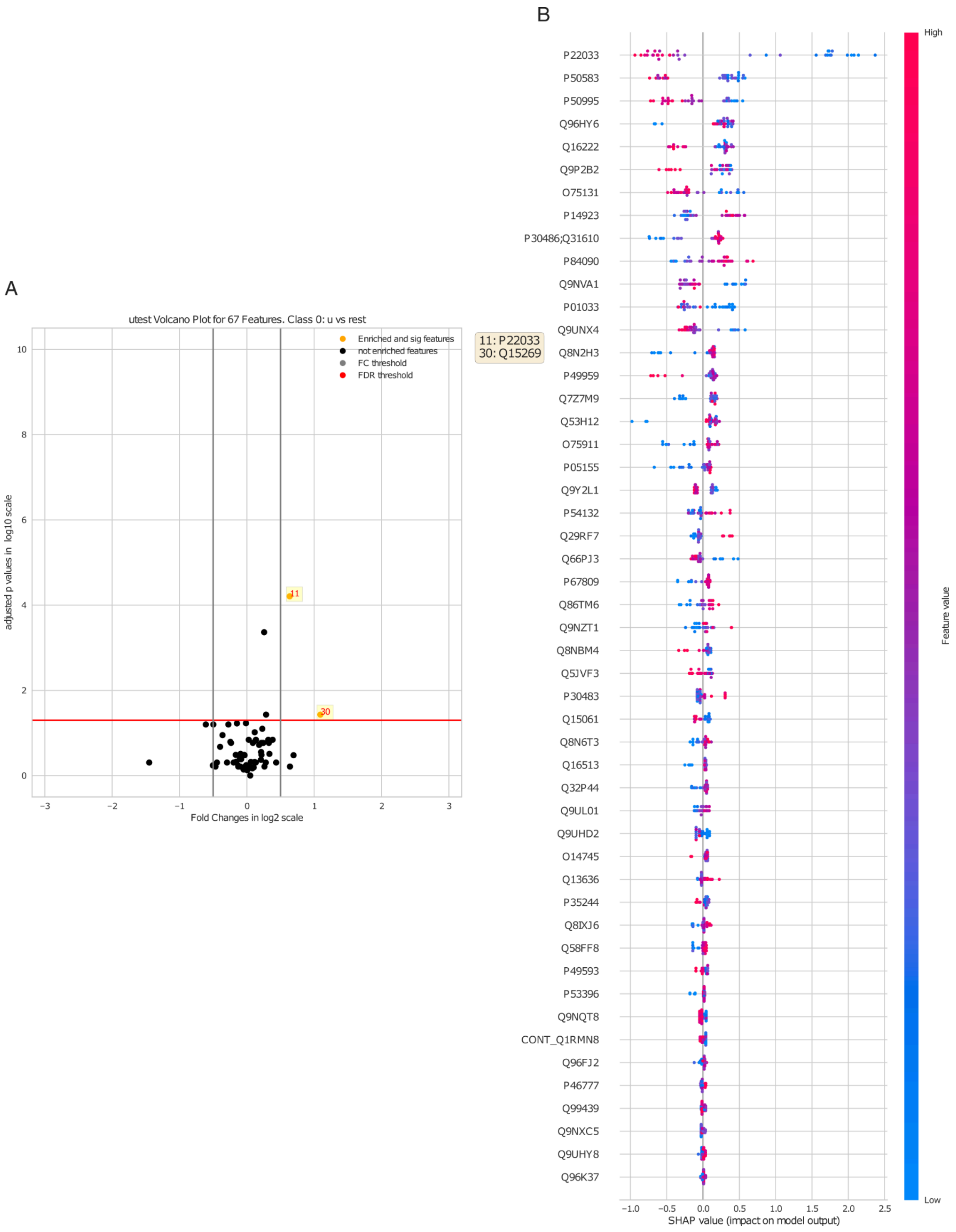


**Figure SF6:** The volcano (SF6A) and SHAP value (SF6B) plots are examples of the output for the 67 MOAgent selected features in the MMA case study on the protein level.

### Correlation Analysis and Feature Distribution:

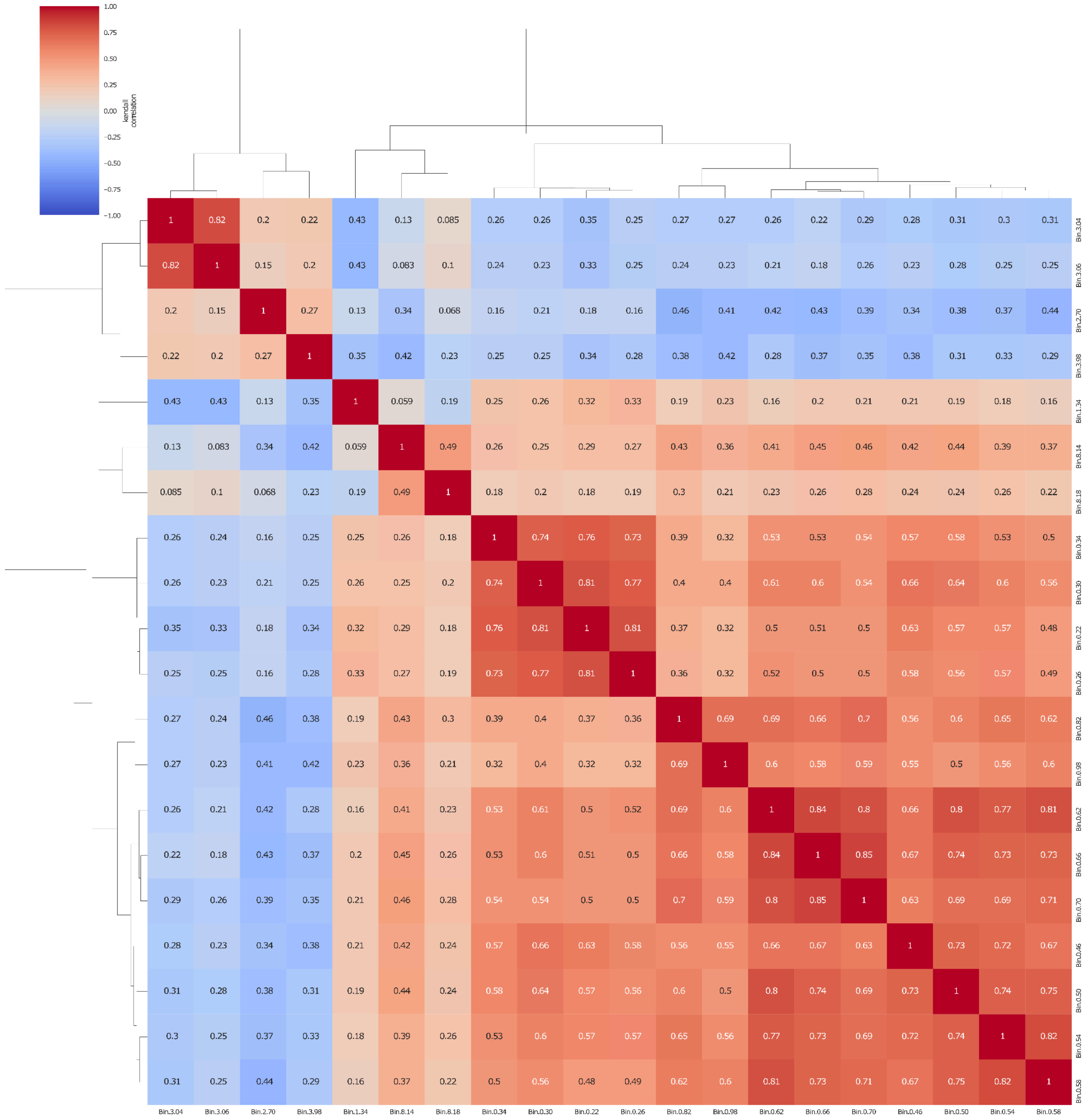


**Figure SF7:** Example correlation heatmap and hierarchical clustering of the features selected by MOAgent in the GN cohort case study.

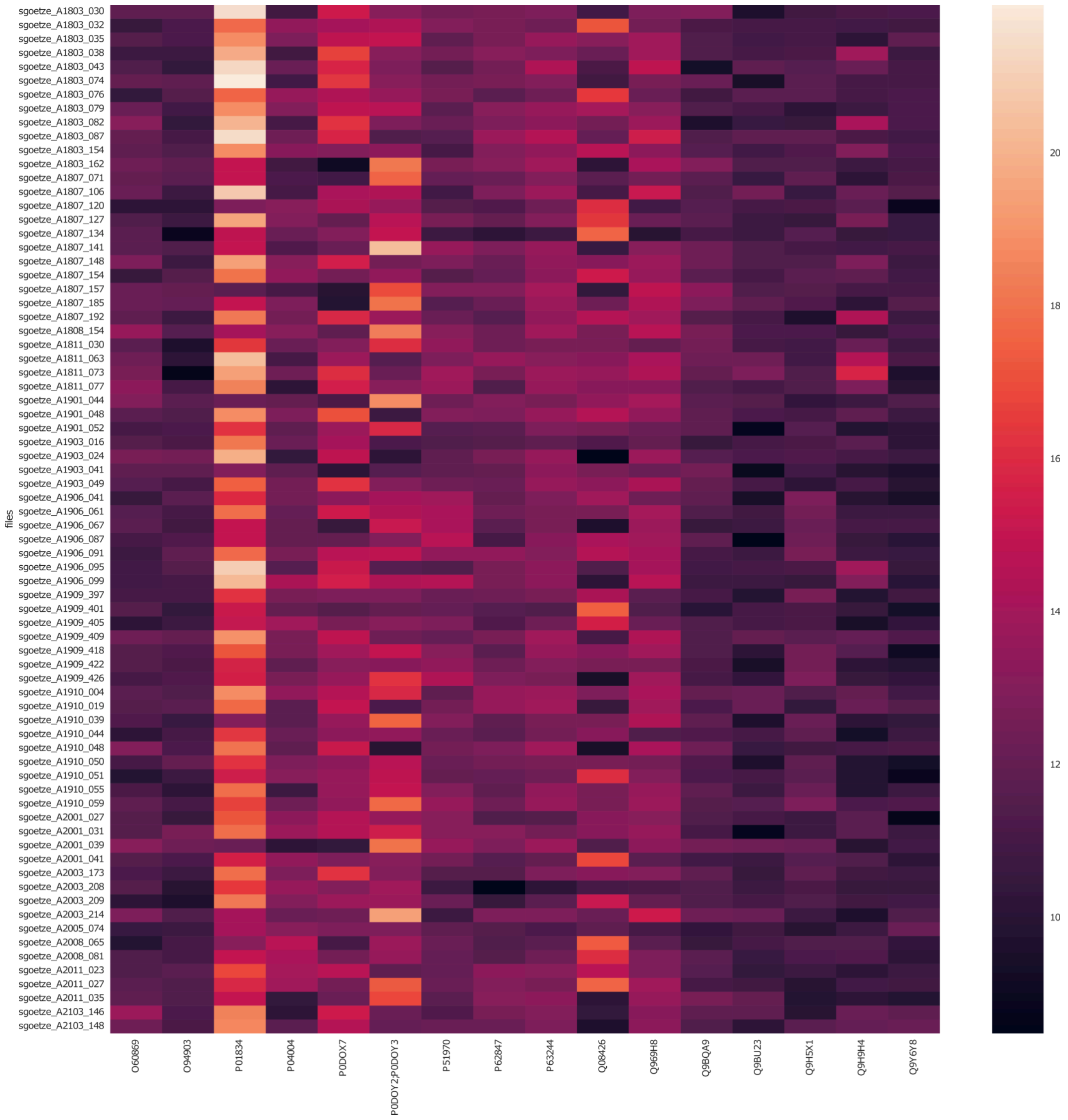

**Figure SF8:** Feature expression heatmap on the protein level of the Kappa-Lambda case study.

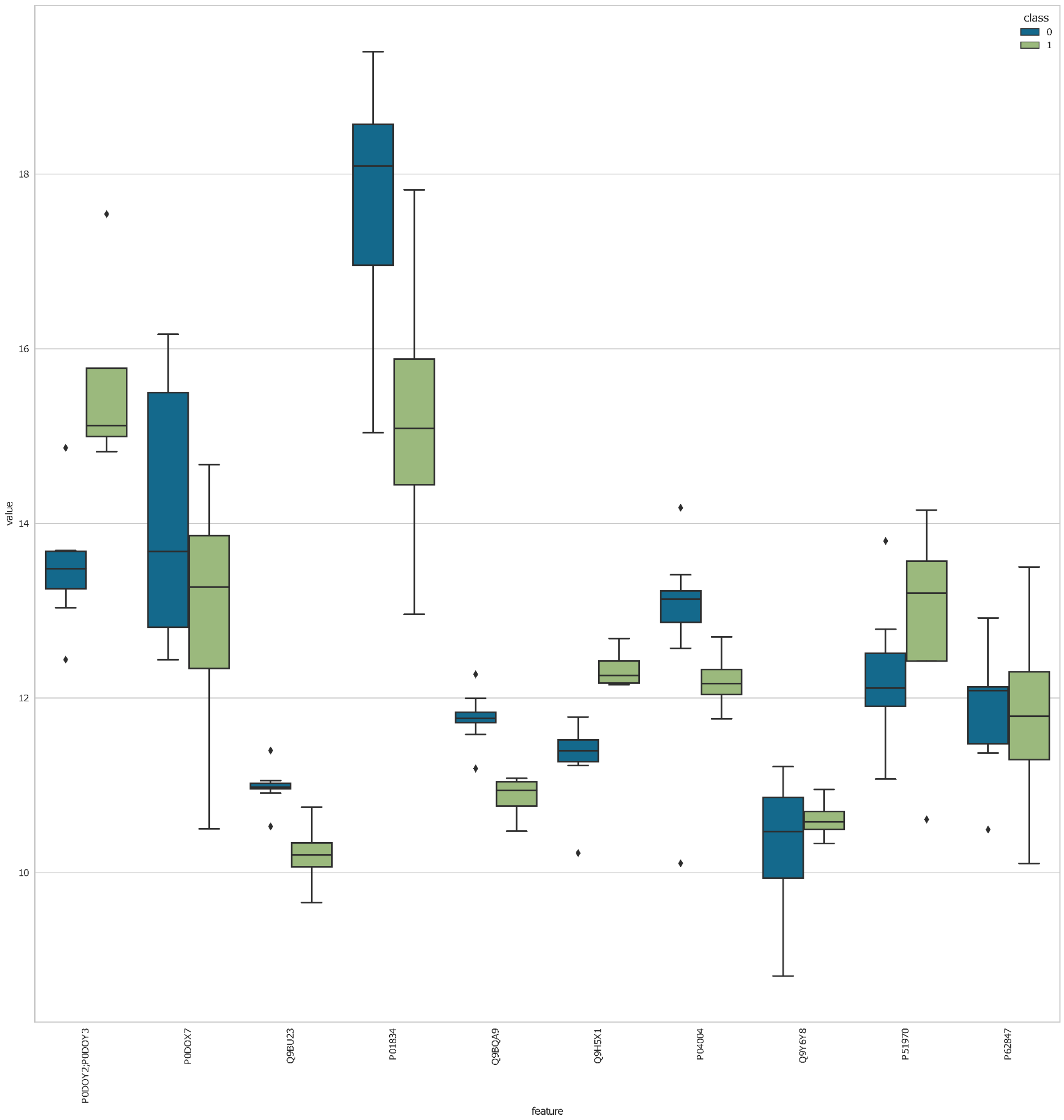


**Figure SF9:** Box plot visualizations of the selected features by MOAgent in the Kappa-Lambda case study on protein level.

### 2. Supplementary Tables

| ID from the original MPN study | Identified by Fragpipe | Selected by MOAgent (\|correlation\| > 0.5, adj. p < 0.01) |
|---|---|---|
| NEK5 (Q6P3R8) | no | no |
| MCM4 (P33991) | yes | no |
| ALPL (P05186) | yes | ALPL (P05186) |
| FH (P07954) | yes | TUFM (P49411) |
| CALR (P27797) | yes | CALR (P27797) |
| STYX (Q8WUJ0) | no | no |
| PSAT1 (Q9Y617) | yes | PSAT1 (Q9Y617) |
| ZNF735 (P0CB33) | no | no |
| COPS8 (Q99627) | yes | RFLNB (Q8N5W9) |
| CD59 (P13987) | yes | CD59 (P13987) |
| NAMPT (P43490) | yes | NAMPT (P43490) |
| ACSL1 (P33121) | yes | CKAP4 (Q07065) |
| S100A9 (P06702) | yes | no |
| MCM7 (P33993) | yes | KDM1A (O60341) |
| RFC2 (P35250) | yes | TAF6 (P49848) |
| POTEJ (P0CG39) | yes | no |
| PSTPIP1 (O43586) | yes | ATP6V1G1 (O75348) |
| PSMB8 (P28062) | yes | ALDH2 (P05091) |
| TPM3 (P06753) | yes | TPM3 (P06753) |
| STOML2 (Q9UJZ1) | yes | UQCRC2 (P22695) |
| CST7 (O76096) | yes | ALG1 (Q9BT22) |

**Table 1:** The table provides an overview of the original 21 most phenotype discriminative proteins found in the original MPN study by Wildschutt et. al. and which of them were found

with the protein identification and quantification platform Fragpipe. The last column provides from MOAgent identified most phenotype discriminative proteins which are correlated (using Kendall’s Tau method) to the original proteins with an absolute correlation value exceeding 0.5 and significance filtered with Benjamini-Hochberg corrected p-values smaller than 0.01.
